# mEos4b photoconversion efficiency depends on laser illumination conditions used in PALM

**DOI:** 10.1101/2022.03.31.486573

**Authors:** Jip Wulffele, Daniel Thedié, Oleksandr Glushonkov, Dominique Bourgeois

## Abstract

Green-to-red photoconvertible fluorescent proteins (PCFPs) are widely employed as markers in photoactivated localization microscopy (PALM). However, their highly complex photophysical behavior complicates their usage. The fact that only a limited fraction of a PCFP ensemble can form the photoconverted state upon near-UV light illumination, termed photoconversion efficiency (PCE), lowers the achievable spatial resolution in PALM and creates undercounting errors in quantitative counting applications. Here, we show that the PCE of mEos4b is not a fixed property of this PCFP, but strongly depends on illumination conditions. Attempts to reduce long-lived blinking in red mEos4b by application of 488 nm light leads to a reduction of the PCE. Furthermore, the PCE of mEos4b strongly depends on the applied 405-nm power density. A refined photophysical model of mEos4b accounts for the observed effects, involving nonlinear green-state photobleaching upon violet light illumination favored by photon absorption by a putative radical dark state.

**TOC GRAPHICS:** 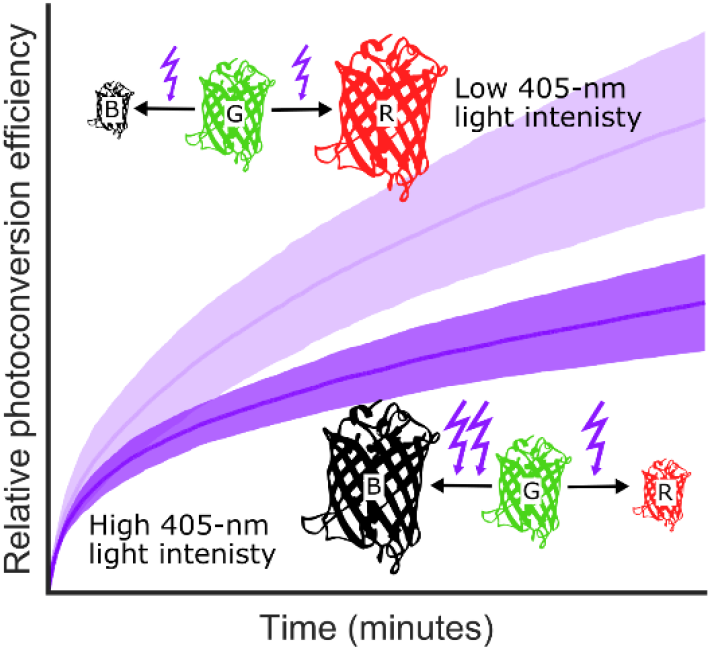

---

For now more than a decade, super-resolution microscopy has provided unprecedented insights into biological processes.^1^ PhotoActivation Localization Microscopy (PALM) is a single-molecule based localization technique that typically takes advantage of so-called phototransformable fluorescent proteins (PTFPs).^2^ Major advantages of using PTFPs are the possibility to work on living cells, the ideal specificity of labeling and the option to label endogenous targets of interest with genome editing techniques.^2^ Green-to-red photoconvertible fluorescent proteins (PCFPs) of the mEos family are often used as PTFP markers in PALM because of their relatively high brightness, high monomeric character, and also because successfully labeled samples can be easily detected by monitoring green state fluorescence prior to PALM data collection.^3^ However, a disadvantage of mEos-based PCFPs is their highly complex photophysical behavior (Supplementary Figure S1).^4–7^

One property essential to the success of PALM experiments using PCFPs is the so-called photoconversion efficiency (PCE),^8,9^ sometimes also referred to as signaling efficiency.^10^ Typically, upon prolonged violet light illumination, green-to-red photoconversion remains incomplete, meaning that from a pool of PCFPs, only a limited fraction can ever be imaged in the red channel of the microscope.^4,9,11,12^ In addition to unsuccessful protein folding or chromophore maturation, incomplete PCE’s can be attributed to possible sub-populations within the PCFP pool unable to photoconvert, rapid photobleaching in the red state before detection can be made, or premature photobleaching in the green state, before photoconversion occurs.^4,9,11,12^ In general, green-to-red photoconversion, the mechanism of which remains incompletely understood,^3^ is in competition with other photophysical pathways that, upon absorption of a violet photon, may lead to reversible dark state formation or photobleaching.^9^

A limited PCE is highly detrimental to PALM experiments. First, it limits the achievable spatial resolution of nanoscopy images due to a reduced apparent labeling efficiency, compromising the high labeling density necessary to satisfy the Nyquist criterium.^8^ Second, it complicates counting by quantitative PALM (qPALM).^12^ In this technique the stoichiometry of protein complexes, typically non-resolvable spatially, can in principle still be determined by counting fluorescence bursts from each complex.^12–16^ However, a limited PCE results in severe undercounting errors. Such errors can in theory be corrected for, based on the expected binomial distribution of measured stoichiometries, but this comes at the cost of increased uncertainty of the extracted values, notably when a distribution of stoichiometries is inherently present in the sample.^12,16^

PCEs of various PCFPs have been reported in several papers, ^9,10,12,13,17,18^ usually relying on the use of biological templates.^14^ The measured values, however, tend to differ widely for a single PCFP, e.g. from ~1%^10^ to ~90%^17^ for mEos2. One hypothesis to account for such variability could be that the employed templates might induce biases due to factors such as template heterogeneity, reduced tumbling of the labels in fixed samples or uncontrolled energy transfer mechanisms between neighboring PCFPs.^11–13^ However, the possibility that PCEs may depend on the employed illumination conditions, notably the applied power densities, has not been put forward. Here, we investigate the effects of 405 and 488-nm light illumination on the PCE of mEos4b, a now popular mEos variant initially designed to withstand chemical fixation.^19^

The study was initially stimulated by the prospect of extending to qPALM applications a blinking reduction strategy that we recently introduced in the context of single-particle-tracking PALM (sptPALM).^5^ In this technique, 488-nm light is used in addition to readout (561-nm) and very weak activating (405-nm, of the order of mW/cm^2^) lights to increase the length of single-molecule tracks through a reduction of fluorescence intermittencies in red mEos4b molecules.^5^ The mechanistic interpretation of the effect is that long off-times in mEos-based PCFPs originate at least in part from 561-nm induced *cis-trans* isomerization of the chromophore, and that back isomerization to the fluorescent *cis* state is efficiently promoted by 488-nm light.^5^ Thus, we wondered whether the same strategy could be used to improve counting accuracy with qPALM, by reducing long-lived blinking expected to be at the origin of enhanced overcounting errors.

We first performed simulations to check whether long-lived off-time reduction effectively improves counting, deliberately ignoring the rich photophysics of mEos4b in its green state.^4,6^ qPALM data were simulated using the photophysical model depicted in Supplementary Figure S2A and Supplementary Table S1, in the absence or the presence of additional 488-nm light. Then, keeping in mind that widely different strategies have been proposed to process qPALM data,^20^ we analyzed the data using either the off-times thresholding method of Lee et al,^15^ or the blinking statistics method of Heilemann et al.^17,16,13,21^ The results, shown in Supplementary Figure S3 and S4, revealed that, whereas the accuracy of stoichiometry retrieval by off times thresholding benefited from long-lived dark state reduction, this was not observed in the case of blinking statistics analysis (Supplementary Note 1). However, importantly, we noticed that slight measurement errors in the blinking propensity (p-value) of mEos4b measured in the monomeric state propagated to very large errors when evaluating complexes of high stoichiometries (Supplementary Figure S4F). Overall, the performed simulations encouraged us to experimentally test the potential benefit of additional 488-nm light for counting purposes.

mEos4b proteins were immobilized in polyacrylamide (PAA) gel and fluorescent traces of photoconverted molecules were recorded using 500 W/cm^2^ 561-nm light under increasing 488-nm light intensities using wide-field illumination. First, we validated that 488-nm illumination reduces the off-time duration of mEos4b in the presence of weak 405-nm laser light (1 W/cm^2^, which can be taken as representative for qPALM experiments)^22^ (Figure 1A). We noticed however that, at similar illumination intensities, the effect was not as pronounced as with 405-nm light (Supplementary Figure S5). The mechanism behind this experimental observation remains to be deciphered, but it could result from the presence of one or several long-lived dark states, in addition to the switched-off protonated *trans* chromophore, which would be reactive to 405-nm but insensitive to-488 nm.

**Figure 1.**
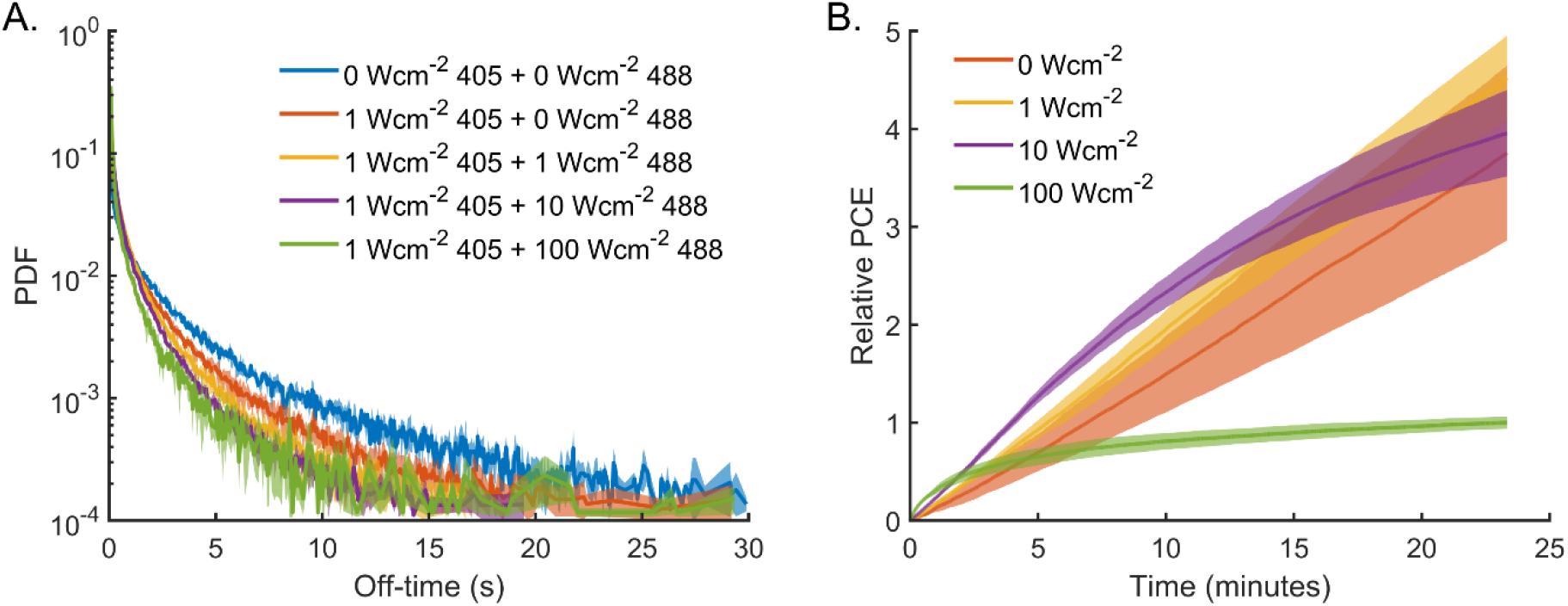
Off-time reduction by 488-nm light is offset by a reduced photoconversion efficiency. Off-time histograms (A) and the relative PCE (B) of mEos4b molecules embedded in PAA under alternating illumination with 500 W/cm^2^ 561-nm light (70 ms), 1 W/cm^2^ 405-nm and 0-100 W/cm^2^ 488-nm light (8.2 ms). Data is normalized to the final number of photoconverted molecules under 100 W/cm^2^ 488 illumination. Data represent mean ± s.d. of ≥ 3 measurements.

Next, we assessed the effect of 488-nm illumination on the PC rate and efficiency. Although absolute PCEs could not be reliably extracted using our experimental set-up (Supplementary Note 4), relative values over time at different illumination intensities could be compared. We thus measured the numbers of photoconverted molecules at the end of experiments using 405-nm, 488-nm and 561-nm light illumination and compared them with those using only 405-nm and 561-nm light. Figure 1B shows that, although low 488-nm light intensity (1 W/cm^2^) had a minimal effect on the PC kinetics within the time window of the experiment, high 488-nm light intensity (100 W/cm^2^) drastically decreased the PCE. Since the bleaching-time and photon budget of photoconverted red molecules were not reduced by 488-nm illumination (Supplementary Figure 8), green state photophysics are likely at the origin of the decrease in PCE. Using simulations with the current model of mEos4b photophysics,^4^ we found that the decrease can be explained by green-state bleaching by the 488-nm light (Supplementary Figure 7). Importantly, the dependence of the PCE on 488-nm light is only observed in the presence of significant 405-nm light (~1 W/cm^2^) (Supplementary Figure S7). This is explained by the fact that in such case photoconversion (mostly driven by 405-nm light) and green-state photobleaching (driven by all lasers) become strongly decoupled. Thus, under very weak 405-nm light (<< 1 W/cm^2^) often used in sptPALM to achieve strong single-molecule sparsity,^5^ there is little penalty in using 488-nm light to reduce red-state intermittencies, although under such circumstances, it has to be kept in mind that the absolute PCE remains very low due to progressive photobleaching by the readout 561-nm light.^4^ Overall, these results show that 488-nm light illumination is not beneficial in qPALM measurements with mEos4b, as the gain in red-state off-time reduction is relatively limited and the PCE becomes compromised in the presence of > ~1 W/cm^2^ 405-nm light.

Intrigued by the finding that 488-nm light drastically reduces the mEos4b PCE in the presence of 405-nm light, we set out to investigate how the intensity of the 405-nm light itself affects the PCE in the absence of 488-nm light. We first hypothesized that the PCE could increase with increasing 405-nm light because 405-nm induced photoconversion and photobleaching rates would evolve in proportion if based on single-photon mechanisms while bleaching of green molecules by the 561-nm laser (ref 4) would be reduced due to shorter acquisition times^4^. To test this, we acquired single-molecule data at varying 405-nm power densities, similar to the 488-nm illumination experiments described above. Whereas low levels of 405-nm light (1 W/cm^2^) increased the PCE as compared to readout photoconversion only, in line with our previous findings,^4^ we found that high 405 intensities (100 W/cm^2^) substantially decreased the PCE despite a higher initial photoconversion rate (Figure 2A). Ensemble measurements where we monitored the cumulative fluorescence of the mEos4b red state under identical light illumination conditions as in the single-molecule case revealed a similar relation between the 405-nm light intensity and the PCE (Supplementary Figure S8). To evaluate whether the decrease in PCE at high 405-nm light intensity could be due to increased bleaching of red molecules, we examined the single-molecule bleaching-time and photon-budget histograms of the red state (Supplementary Figure S9). These histograms reveal that red-state bleaching was not significantly increased when raising the 405-nm light power density, suggesting that, again, green-state photophysics are at the origin of the decreased PCE under increased 405-nm light intensities.

**Figure 2.**
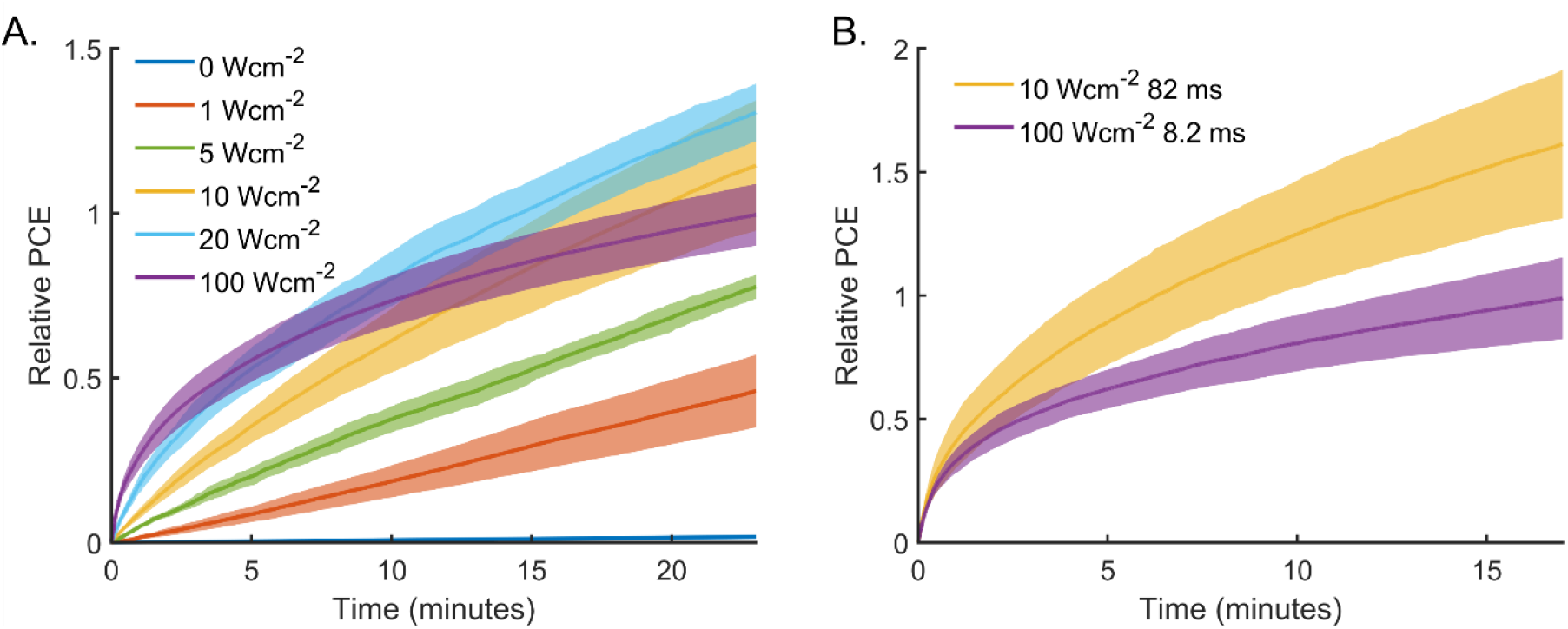
High 405-nm illumination intensities reduce the photoconversion efficiency of mEos4b under PALM imaging conditions. A) Relative PCE of molecules of mEos4b molecules embedded in PAA under alternating illumination with 500 W/cm^2^ 561-nm light (70 ms) and 1-100 W/cm^2^ 405-nm light (8.2 ms). B) Relative number of photoconverted molecules of mEos4b molecules embedded in PAA under alternating illumination with 500 W/cm^2^ 561-nm light (70 ms) and 10 or 100 W/cm^2^ 405-nm light for 82 and 8.2 ms, respectively. Data is normalized to the final number of photoconverted molecules under 100 W/cm^2^ 405 illumination. Data represent mean ± s.d. of ≥ 3 measurements.

In contrast to the case of 488-nm light, the dependence of the mEos4b PCE on 405-nm light cannot be explained by the current photophysical model of this PCFP (Supplementary Figure S7).^4^ To explain the decrease of the PCE at high 405-nm light intensity, 405-nm photons must bleach the green state in a non-linear manner, that is, bleaching must require the subsequent absorption of two (or more) photons. To verify this hypothesis, we compared the PCEs under 10 W/cm^2^ and 100 W/cm^2^ 405-nm light while keeping the integrated illumination dose constant by adjusting the exposure time (82 ms for 10 W/cm^2^, 8.2 ms for 100 W/cm^2^). As expected, application of short exposures of 100 W/cm^2^ 405-nm light decreased the mEos4b PCE as compared to longer exposures to 10 W/cm^2^, while this did not affect the red state photophysics (Figure 2B, Supplementary Figure S10).

Next, we aimed to identify the photophysical state(s) of green mEos4b involved in non-linear bleaching by 405-nm light. Violet light is predominantly absorbed by the protonated states of the green chromophore,^6^ which makes the *cis*-protonated and *trans*-protonated (i.e. off-switched) states likely starting points for the mechanism at play. The *trans*-protonated state can be populated by intense 561-nm illumination,^4^ as used in PALM measurements to reach single-molecule sensitivity, which made us wonder whether 405-nm light also induced non-linear bleaching in the absence of 561-nm light. To test this, we monitored the progression of ensemble mEos4b red fluorescence under 1, 10 and 100 W/cm^2^ 405-nm illumination and weak exposure to 561-nm light (8 W/cm^2^) (Figure 3A). Sun et al^7^ reported that the ensemble red fluorescence intensity of mEos3.2 under constant 405-nm and 561-nm illumination follows a 3-state model: G -> R -> B. Such model implies that the level of the maximum recorded red fluorescence increases with the applied 405-nm light intensity as bleaching by the 561-nm laser lowers the peak height at low 405-nm intensities. However, the experimentally measured maximum intensity under 100 W/cm^2^ 405-nm light reached a significantly lower value as compared to illumination with 1 and 10 W/cm^2^ 405-nm light (Figure 3A, Supplementary Note 3), suggesting that the model of Sun et al is insufficient to account for our data. To further validate this finding we monitored the ensemble red-state fluorescence under illumination with either 10 or 100 W/cm^2^ 405-nm light while keeping the integrated dose of 405-nm light constant. The data again revealed that high-intensity 405-nm light decreased the red fluorescence maximum intensity (Figure 3B). Altogether, these ensemble measurements are consistent with the notion that 405-nm light induces non-linear photobleaching of the mEos4b green state independently of the presence of additional 561-nm light. We speculate that upon absorption of a 405-nm photon in either the *cis*-neutral or *trans*-neutral chromophore state, an intermediate state forms with a minimum lifetime in the tens of milliseconds range (Figure 4, Supplementary Note 5). Such lifetime appears too long to attribute the intermediate to the triplet state.^23^ However, a radical state, similar to the one that we pinpointed in an earlier X-ray-based study of IrisFP (a PCFP derived from EosFP), could be invoked^24^. This intermediate would thermally relax to the fluorescence state or lead to photobleaching upon absorption of violet light. The precise mechanism of non-linear bleaching of PCFPs by 405-nm light will now need further dedicated studies.

**Figure 3.**
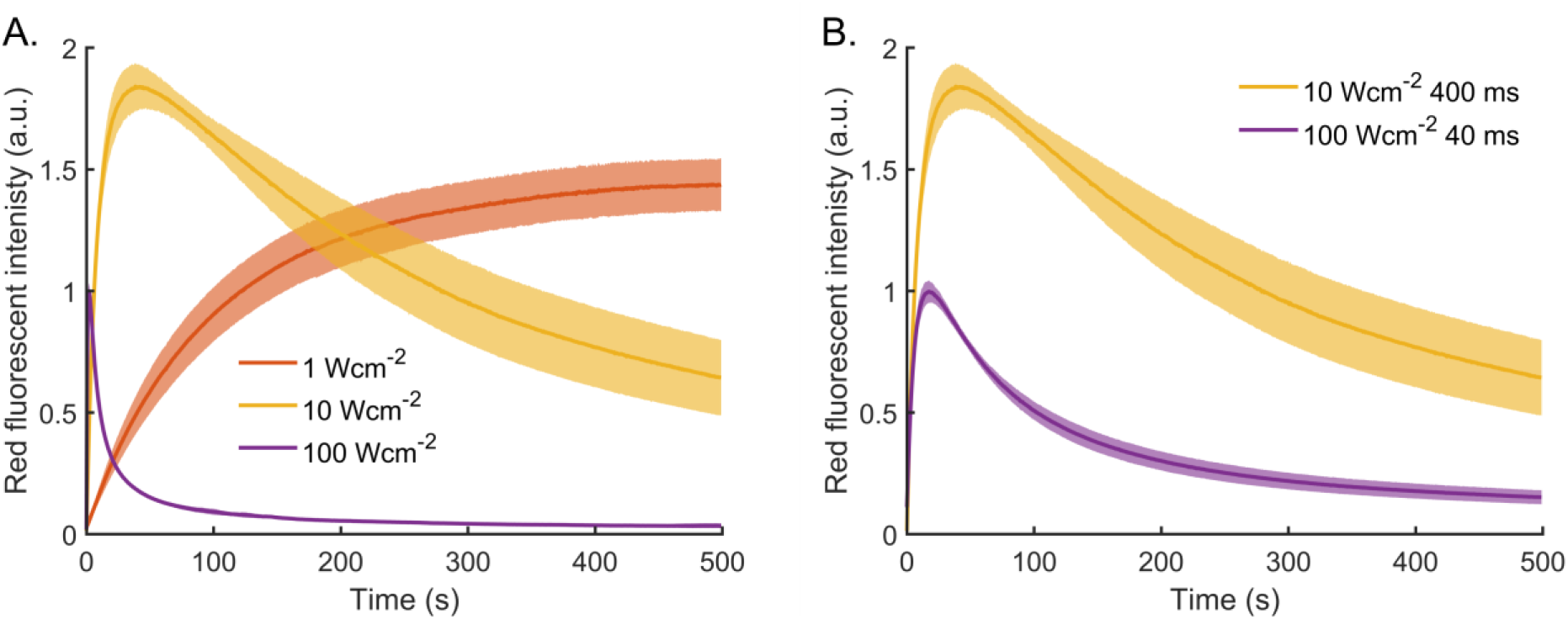
High 405-nm light illumination intensities reduce the photoconversion efficiency of mEos4b in the absence of 561-nm light illumination. Ensemble fluorescence time traces of mEos4b molecules embedded in PAA. The red fluorescent intensity was measured every 500 ms by illumination with 8 W/cm^2^ 561-nm light for 20 ms. A) Samples were exposed to 400 ms pulses of 405-nm light (1-100 W/cm^2^). B) Samples were exposed to 40 and 400 ms pulses of 100 and 10 W/cm^2^ 405-nm respectively. Data represent mean ± s.d. of ≥ 3 measurements.

**Figure 4.**
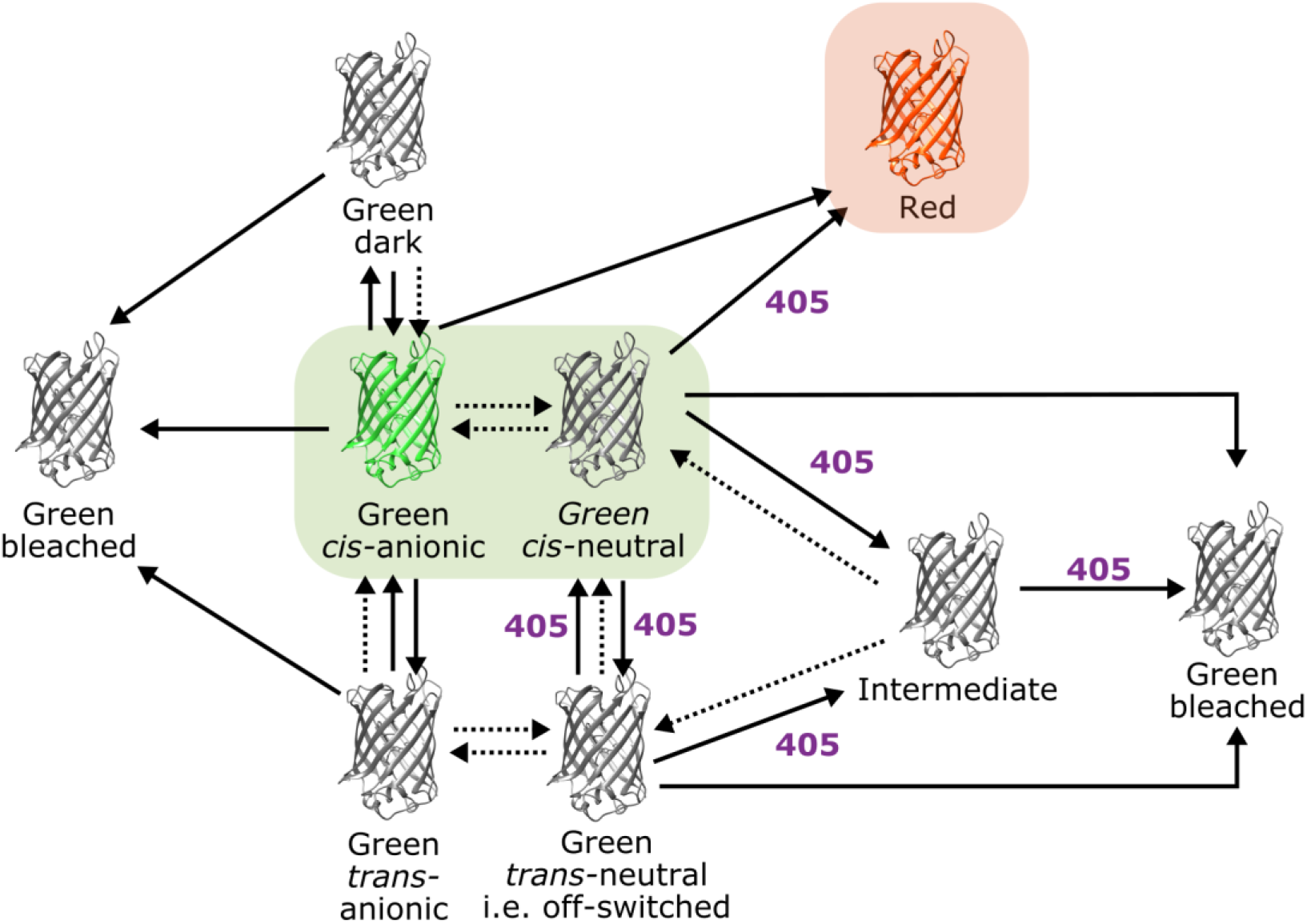
Tentative updated model of mEos4b green-state photophysics. An intermediate state, downstream of the neutral green chromophore, thermally recovers or absorbs a second 405-nm photon leading to nonlinear bleaching at high 405-nm power densities. Single-photon bleaching may occur from every light absorbing state. Solid arrows indicate photo-induced transitions; dotted arrows indicate thermal transitions or pH-dependent equilibria.

In conclusion, our results demonstrate that the photoconversion efficiency of mEos4b strongly depends on the employed illumination conditions. In the presence of 405-nm light the PCE of mEos4b decreases with increasing 488-nm illumination because 488-nm light significantly increases the overall green-state bleaching rate while the overall photoconversion rate remains governed by the applied 405-nm light. Most importantly, high 405-nm light intensities decrease the PCE of mEos4b due to non-linear bleaching of the green state, possibly through subsequent photon absorptions by a neutral state of mEos4b and a short-lived radical state. Accordingly, an updated model of mEos4b photophysics is presented in Figure 4, of use for the future design of optimized illumination schemes. We also observed nonlinear photobleaching by 405-nm light in mEos3.2 (Supplementary Figure 11) and we anticipate that the described mechanism applies to all commonly used mEos variants, including mEos2 and mEos3.1. In the future, it will be interesting to investigate the effect of pH, oxygen removal, addition of triplet state quenchers, reducers or oxidizers, on the nonlinear photobleaching phenomenon.^25,26^ Other types of green-to-red photoconvertible fluorescent proteins such as priming PCFPs^27^ may behave differently. A high PCE is essential to ensure sufficient labeling density in PALM and accurate molecular counting in qPALM. Whereas the use of very low 405-nm light (<< 1 W/cm^2^) should be avoided whenever possible (depending on single-molecule sparsity) due to accumulating green-state photobleaching by readout 561-nm light, high 405-nm light (> ~20 W/cm^2^) is expected to also yield sub-optimal results in the following circumstances: *(i)* in high-speed PALM;^28^ *(ii)* whenever the photoconversion light is progressively ramped-up in PALM or qPALM; or *(iii)* whenever short and intense photoconversion light pulses (e.g. in between recorded frames) are used instead of longer and weaker pulses. Finally, this work highlights the complexity and, yet, key importance of green-state photophysics in photoconvertible fluorescent proteins.

## ASSOCIATED CONTENT

The following files are available free of charge.

Supplementary information (PDF)

## AUTHOR INFORMATION

JW & DB conceived of the project; JW & DT performed experiments and data analysis; JW & DB performed simulations; OG & DT developed data acquisition schemes; JW & DB wrote the paper with contributions from all authors.

The authors declare no competing financial interests.

## ACKNOWLEDGMENT

We thank Ninon Zala, Virgile Adam, Joanna Timmins and Salvatore De Bonis for PCFP’s production and purification, and Jean-Philippe Kleman for assistance with the M4D imaging platform. This work was supported by the Agence Nationale de la Recherche (grant no. ANR-17-CE11-0047-01 and ANR-20-CE11-0013-01 to D.B. and ANR-16-CE11-0016-01) and used the M4D imaging platform of the Grenoble Instruct-ERIC Center (ISBG: UMS 3518 CNRS-CEA-UGA-EMBL) with support from FRISBI (grant no. ANR-10-INBS-05-02) and GRAL, a project of the Université Grenoble Alpes graduate school (Ecoles Universitaires de Recherche) CBH-EUR-GS (ANR-17-EURE-0003) within the Grenoble Partnership for Structural Biology (PSB). JW acknowledges funding by the GRAL Labex.

## REFERENCES

(1) Lelek, M.; Gyparaki, M. T.; Beliu, G.; Schueder, F.; Griffié, J.; Manley, S.; Jungmann, R.; Sauer, M.; Lakadamyali, M.; Zimmer, C. Single-Molecule Localization Microscopy. Nat. Rev. Methods Primer 2021, 1 (1), 1–27. https://doi.org/10.1038/s43586-021-00038-x.

(2) Shcherbakova, D. M.; Sengupta, P.; Lippincott-Schwartz, J.; Verkhusha, V. V. Photocontrollable Fluorescent Proteins for Superresolution Imaging. Annu. Rev. Biophys. 2014, 43, 303–329. https://doi.org/10.1146/annurev-biophys-051013-022836.

(3) Nienhaus, K.; Ulrich Nienhaus, G. Fluorescent Proteins of the EosFP Clade: Intriguing Marker Tools with Multiple Photoactivation Modes for Advanced Microscopy. RSC Chem. Biol. 2021, 2 (3), 796–814. https://doi.org/10.1039/D1CB00014D.

(4) Thédié, D.; Berardozzi, R.; Adam, V.; Bourgeois, D. Photoswitching of Green MEos2 by Intense 561 Nm Light Perturbs Efficient Green-to-Red Photoconversion in Localization Microscopy. J. Phys. Chem. Lett. 2017, 8 (18), 4424–4430. https://doi.org/10.1021/acs.jpclett.7b01701.

(5) De Zitter, E.; Thédié, D.; Mönkemöller, V.; Hugelier, S.; Beaudouin, J.; Adam, V.; Byrdin, M.; Van Meervelt, L.; Dedecker, P.; Bourgeois, D. Mechanistic Investigation of MEos4b Reveals a Strategy to Reduce Track Interruptions in SptPALM. Nat. Methods 2019, 16 (8), 707–710. https://doi.org/10.1038/s41592-019-0462-3.

(6) De Zitter, E.; Ridard, J.; Thédié, D.; Adam, V.; Lévy, B.; Byrdin, M.; Gotthard, G.; Van Meervelt, L.; Dedecker, P.; Demachy, I.; Bourgeois, D. Mechanistic Investigations of Green MEos4b Reveal a Dynamic Long-Lived Dark State. J. Am. Chem. Soc. 2020, 142 (25), 10978–10988. https://doi.org/10.1021/jacs.0c01880.

(7) Sun, M.; Hu, K.; Bewersdorf, J.; Pollard, T. D. Sample Preparation and Imaging Conditions Affect MEos3.2 Photophysics in Fission Yeast Cells. Biophys. J. 2021, 120 (1), 21–34. https://doi.org/10.1016/j.bpj.2020.11.006.

(8) Diekmann, R.; Kahnwald, M.; Schoenit, A.; Deschamps, J.; Matti, U.; Ries, J. Optimizing Imaging Speed and Excitation Intensity for Single-Molecule Localization Microscopy. Nat. Methods 2020, 17 (9), 909–912. https://doi.org/10.1038/s41592-020-0918-5.

(9) Durisic, N.; Laparra-Cuervo, L.; Sandoval-Álvarez, Á.; Borbely, J. S.; Lakadamyali, M. Single-Molecule Evaluation of Fluorescent Protein Photoactivation Efficiency Using an in Vivo Nanotemplate. Nat. Methods 2014, 11 (2), 156–162. https://doi.org/10.1038/nmeth.2784.

(10) Wang, S.; Moffitt, J. R.; Dempsey, G. T.; Xie, X. S.; Zhuang, X. Characterization and Development of Photoactivatable Fluorescent Proteins for Single-Molecule–Based Superresolution Imaging. Proc. Natl. Acad. Sci. 2014, 111 (23), 8452–8457. https://doi.org/10.1073/pnas.1406593111.

(11) Tao, A.; Zhang, R.; Yuan, J. Characterization of Photophysical Properties of Photoactivatable Fluorescent Proteins for Super-Resolution Microscopy. J. Phys. Chem. B 2020, 124 (10), 1892–1897. https://doi.org/10.1021/acs.jpcb.9b11028.

(12) Puchner, E. M.; Walter, J. M.; Kasper, R.; Huang, B.; Lim, W. A. Counting Molecules in Single Organelles with Superresolution Microscopy Allows Tracking of the Endosome Maturation Trajectory. Proc. Natl. Acad. Sci. 2013, 110 (40), 16015–16020. https://doi.org/10.1073/pnas.1309676110.

(13) Baldering, T. N.; Dietz, M. S.; Gatterdam, K.; Karathanasis, C.; Wieneke, R.; Tampé, R.; Heilemann, M. Synthetic and Genetic Dimers as Quantification Ruler for Single-Molecule Counting with PALM. Mol. Biol. Cell 2019, 30 (12), 1369–1376. https://doi.org/10.1091/mbc.E18-10-0661.

(14) Finan, K.; Raulf, A.; Heilemann, M. A Set of Homo-Oligomeric Standards Allows Accurate Protein Counting. Angew. Chem. Int. Ed. 2015, 54 (41), 12049–12052. https://doi.org/10.1002/anie.201505664.

(15) Lee, S.-H.; Shin, J. Y.; Lee, A.; Bustamante, C. Counting Single Photoactivatable Fluorescent Molecules by Photoactivated Localization Microscopy (PALM). Proc. Natl. Acad. Sci. 2012, 109 (43), 17436–17441. https://doi.org/10.1073/pnas.1215175109.

(16) Baldering, T. N.; Bullerjahn, J. T.; Hummer, G.; Heilemann, M.; Malkusch, S. Molecule Counts in Complex Oligomers with Single-Molecule Localization Microscopy. J. Phys. Appl. Phys. 2019, 52 (47), 474002. https://doi.org/10.1088/1361-6463/ab3b65.

(17) Fricke, F.; Beaudouin, J.; Eils, R.; Heilemann, M. One, Two or Three? Probing the Stoichiometry of Membrane Proteins by Single-Molecule Localization Microscopy. Sci. Rep. 2015, 5 (1), 14072. https://doi.org/10.1038/srep14072.

(18) Nan, X.; Collisson, E. A.; Lewis, S.; Huang, J.; Tamgüney, T. M.; Liphardt, J. T.; McCormick, F.; Gray, J. W.; Chu, S. Single-Molecule Superresolution Imaging Allows Quantitative Analysis of RAF Multimer Formation and Signaling. Proc. Natl. Acad. Sci. 2013, 110 (46), 18519–18524. https://doi.org/10.1073/pnas.1318188110.

(19) Paez Segala, M. G.; Sun, M. G.; Shtengel, G.; Viswanathan, S.; Baird, M. A.; Macklin, J. J.; Patel, R.; Allen, J. R.; Howe, E. S.; Piszczek, G.; Hess, H. F.; Davidson, M. W.; Wang, Y.; Looger, L. L. Fixation-Resistant Photoactivatable Fluorescent Proteins for Correlative Light and Electron Microscopy. Nat. Methods 2015, 12 (3), 215–218. https://doi.org/10.1038/nmeth.3225.

(20) Khater, I. M.; Nabi, I. R.; Hamarneh, G. A Review of Super-Resolution Single-Molecule Localization Microscopy Cluster Analysis and Quantification Methods. Patterns 2020, 1 (3), 100038. https://doi.org/10.1016/j.patter.2020.100038.

(21) Baldering, T. N.; Karathanasis, C.; Harwardt, M.-L. I. E.; Freund, P.; Meurer, M.; Rahm, J. V.; Knop, M.; Dietz, M. S.; Heilemann, M. CRISPR/Cas12a-Mediated Labeling of MET Receptor Enables Quantitative Single-Molecule Imaging of Endogenous Protein Organization and Dynamics. iScience 2021, 24 (1), 101895. https://doi.org/10.1016/j.isci.2020.101895.

(22) Fricke, F.; Beaudouin, J.; Malkusch, S.; Eils, R.; Heilemann, M. Quantitative Single-Molecule Localization Microscopy (QSMLM) of Membrane Proteins Based on Kinetic Analysis of Fluorophore Blinking Cycles. In Super-Resolution Microscopy: Methods and Protocols; Erfle, H., Ed.; Methods in Molecular Biology; Springer: New York, NY, 2017; pp 115–126. https://doi.org/10.1007/978-1-4939-7265-4_10.

(23) Byrdin, M.; Duan, C.; Bourgeois, D.; Brettel, K. A Long-Lived Triplet State Is the Entrance Gateway to Oxidative Photochemistry in Green Fluorescent Proteins. J. Am. Chem. Soc. 2018, 140 (8), 2897–2905. https://doi.org/10.1021/jacs.7b12755.

(24) Roy, A.; Field, M. J.; Adam, V.; Bourgeois, D. The Nature of Transient Dark States in a Photoactivatable Fluorescent Protein. J. Am. Chem. Soc. 2011, 133 (46), 18586–18589. https://doi.org/10.1021/ja2085355.

(25) Henrikus, S. S.; Tassis, K.; Zhang, L.; van der Velde, J. H. M.; Gebhardt, C.; Herrmann, A.; Jung, G.; Cordes, T. Characterization of Fluorescent Proteins with Intramolecular Photostabilization**. ChemBioChem 2021, 22 (23), 3283–3291. https://doi.org/10.1002/cbic.202100276.

(26) Jusuk, I.; Vietz, C.; Raab, M.; Dammeyer, T.; Tinnefeld, P. Super-Resolution Imaging Conditions for Enhanced Yellow Fluorescent Protein (EYFP) Demonstrated on DNA Origami Nanorulers. Sci. Rep. 2015, 5 (1), 14075. https://doi.org/10.1038/srep14075.

(27) Turkowyd, B.; Balinovic, A.; Virant, D.; Carnero, H. G. G.; Caldana, F.; Endesfelder, M.; Bourgeois, D.; Endesfelder, U. A General Mechanism of Photoconversion of Green-to-Red Fluorescent Proteins Based on Blue and Infrared Light Reduces Phototoxicity in Live-Cell Single-Molecule Imaging. Angew. Chem. Int. Ed Engl. 2017, 56 (38), 11634–11639. https://doi.org/10.1002/anie.201702870.

(28) Development of ultrafast camera-based imaging of single fluorescent molecules and live-cell PALM | bioRxiv https://www.biorxiv.org/content/10.1101/2021.10.26.465864v2 (accessed 2022 -02 -22).

