## Supplementary information for "mEos4b photoconversion efficiency depends on laser illumination conditions used in PALM"

### Material and Methods

#### Purification of mEos4b and sample preparation

mEos4b was purified according to the protocol described in[1]. The purified protein was diluted in 750 mM Tris buffer pH 8 containing 15% Acrylamide/Bis-acrilamide (29:1, Invitrogen) to nanomolar or micromolar concentration for single-molecule and ensemble experiments, respectively. For single-molecule experiments, nanodiamonds (12 ug/ml final concentration, NDN V100nmHi10mL, Adamas Nanotechnologies) were added as fiducials. Next, polymerization was initiated by addition of 1% TEMED (Thermo Scientific) and 0.1% ammonium persulfate (Sigma), after which the gel was evenly spread between 2 coverslips to a thin layer of approximately 10 $\mu$m thickness. The gel was left to polymerize at room temperature for several minutes.

#### Fluorescence microscopy experiments

Ensemble and single-molecule data were acquired on a home-built PALM/STORM set-up based on an Olympus IX81 inverted microscope equipped with a x100 1.49 NA oil-immersion apochromatic objective lens (Olympus). Widefield illumination was achieved by focusing the diode-pumped solid state 405-nm (CrystaLaser), 488-nm (Spectra Physics) and 561-nm (Cobolt Jena) laser beams to the back focal plane of the objective. Intensities of laser illuminations at the sample were tuned by an acousto-optical tunable filter (AOTF, AA Opto Electronic). Fluorescence images were acquired with an Evolve 512 back-illuminated EMCCD camera (Photometrics) controlled by the Metamorph software (Molecular Devices).

##### *Single-molecule data collection*

Single-molecule data were collected with 561-nm light illumination (500W/cm$^2$) using an exposure time of 70 ms. All frames were interleaved with 12 ms time periods during which the sample was exposed to 405-nm and/or 488-nm light illumination (0-100W/cm$^2$ for 8.2 ms).

To evaluate nonlinear photobleaching by 405-nm illumination, the time separating subsequent frames was increased to 82 ms, during which the sample was exposed to 10 or 100W/cm$^2$ 405-nm light for 82 and 8.2 ms respectively.

*Ensemble fluorescence measurements*

Ensemble-level measurements were recorded using an exposure time of 20 ms and 8 W/cm² 561-nm light illumination to readout the red fluorescence. Frames were separated by 480 ms, during which the sample was exposed to 405-nm light illumination (0-100W/cm² for 40 or 400 ms). Alternatively, ensemble-level measurements were recorded under the same conditions as the single molecule data (Supplementary Figure 8).

### Data processing

*Single-molecule data analysis*

Single molecules were localized using the Thunderstorm plugin in ImageJ[2] and drift correction was performed using the nanodiamonds as fiducials. The localizations were clustered to reconstitute the fluorescence time-traces of mEos4b monomers using custom-written Matlab scripts based on the *k-means* clustering algorithm and a maximum allowed off-time duration of 30 seconds.[3] Spurious localizations were removed based on photon budget and localization precision while a fraction of clusters were filtered out based on number of localizations and ellipticity. Next, blinking characteristics (on-times, off-times, bleaching-times, numbers of blinks and photon budgets) were extracted from the fluorescent time-traces and plotted as histograms.

To estimate the relative number of photoconverted molecules under the different illumination conditions tested, the cumulative numbers of photoconverted molecules over time were extracted from the clustered data. To correct for concentration differences between samples, a dataset under 10W/cm² 405-nm light illumination was acquired as a reference on each sample.

*Ensemble data analysis*

Fluorescent time evolutions were extracted from a small area in which the laser power densities was considered homogeneous, and background fluorescence was subtracted. To evaluate the effect of the 405-nm intensity under PALM conditions, the cumulative red fluorescence intensity was calculated from the ensemble data acquired under PALM imaging conditions.

### Simulations

*PALM data sets of mEos4b monomers*

Single-molecule PALM data of mEos4b monomers were simulated using an in house simulation software written in Matlab, able to incorporate the photophysical properties of PCFPs.[3,4] The used photophysical model is described in Supplementary Figure 2 and the employed rates and quantum yields are shown in Supplementary Table 1. No green-state photophysics were needed in those simulations. The following parameters were used: 70 ms frame time, 12 ms in between frames, 500W/cm² 561-nm power density, 1W/cm² 405-nm power density and 0-10W/cm² 488-nm power density for reduced off-times. For each illumination condition, over 20000 mEos4b molecules were simulated.

*Counting analysis*

The simulated data were localized and clustered as described for the experimental PALM data. The fluorescence time traces of 2000 virtual oligomers of various stoichiometries (dimers, tetramers, octamers and 16-mers) were reconstituted by merging the time traces of randomly selected monomers, eventually applying the binomial low to simulate the effect of a reduced PCE. The reconstitute oligomers were then subjected to the two counting methods developed Lee et al (2012) and by Fricke et al (2015).[5,6]

*Effect of green-state photobleaching*

To assess the relative effects of the 405-nm, 488-nm and 561-nm lasers for photoconversion and green-state photobleaching, similar simulations were performed that incorporate green-state photophysics as described in Supplementary Figure S1 and Supplementary Table S2.

### Supplementary Figures

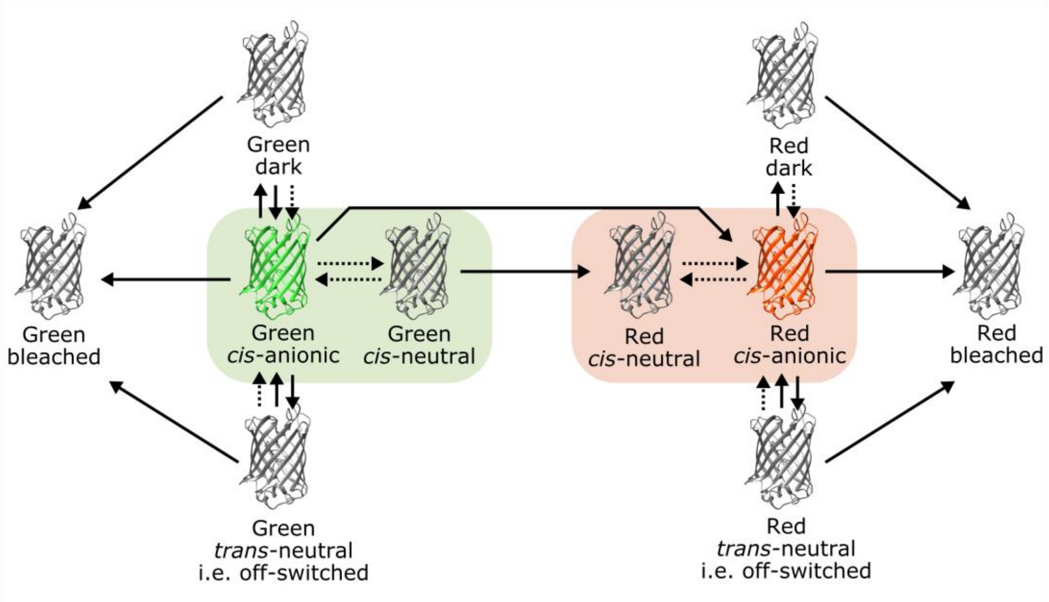


**Supplementary Figure S1. Current photophysical model of mEos4b as proposed by Thedie et al.**[7] Solid arrows indicate photo-induced transitions; dotted arrows indicate thermal transitions or pH-dependent equilibria.

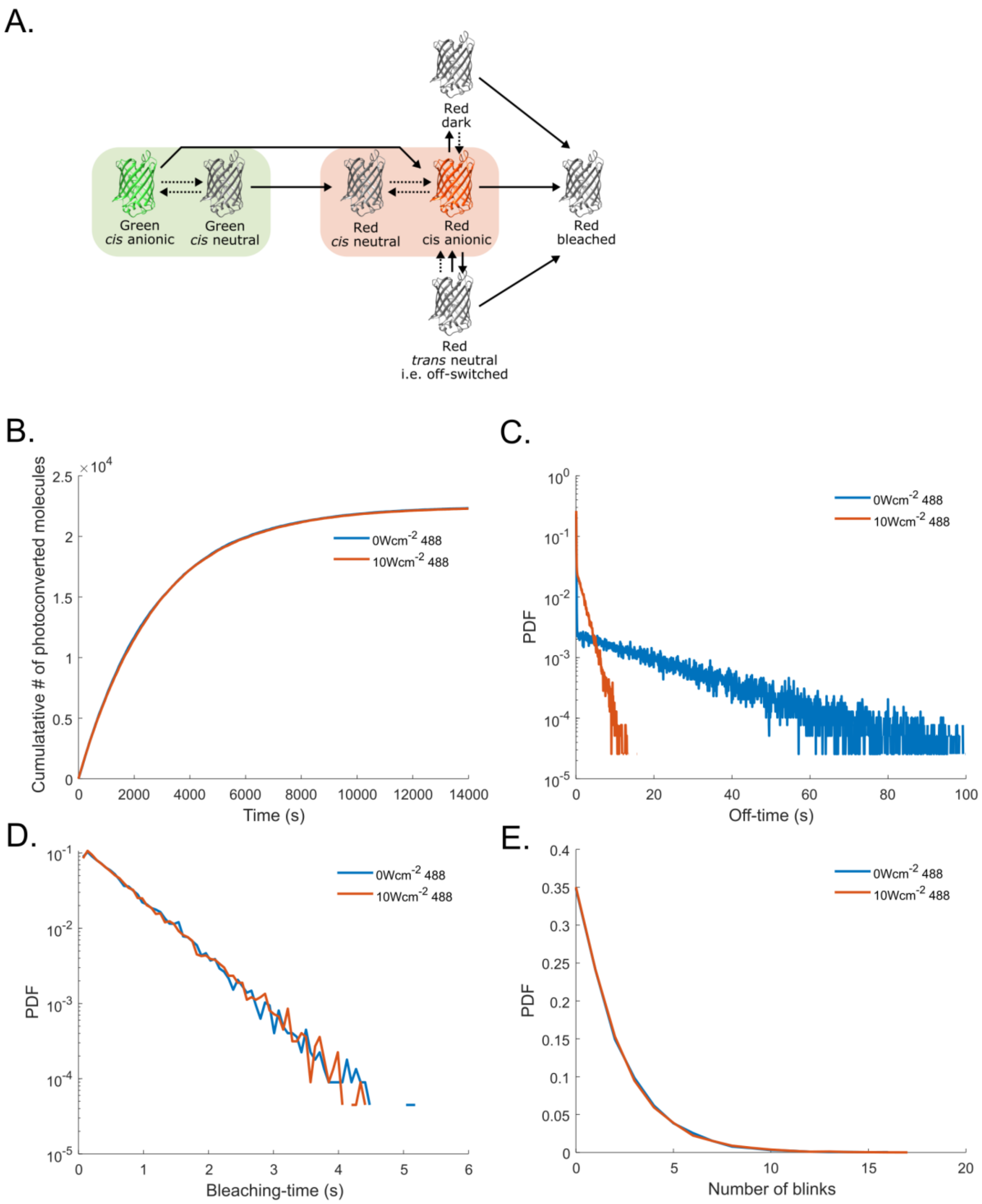


**Supplementary Figure S2. mEos4b simulations for counting analysis.** PALM imaging of mEos4b was simulated under illumination with 500W/cm$^2$ 561-nm and 1W/cm$^2$ 405-nm light, typical for qPALM, with and without the addition of 10W/cm$^2$ 488-nm light to reduce red-state off-time durations. A) Photophysical model of mEos4b used for PALM simulations. Green-state photophysics were omitted to be able to focus solely on the effect of red-state off-time reduction on molecular counting. Solid arrows indicate photo-induced transitions; dotted arrows indicate thermal transitions or pH-dependent equilibria. See Supplementary Table 1 for the corresponding phototransformation quantum yields and thermal exchange rates. B) Using the simplified photophysical model of mEos4b, addition of 488-nm light minimally affect the PC kinetics due to additional PC from the cis-anionic green chromophore. C) Addition of 488-nm light illumination significantly decreases the red-state off-time durations of simulated mEos4b molecules. D-E) Addition of 10W/cm$^2$ 488-nm light illumination minimally affects the red-state nBlink (D) and bleaching-time (E) histograms of simulated mEos4b molecules.

**Supplementary Table S1. Phototransformation quantum yields and thermal exchange rates used for the PALM simulations of mEos4b.**

| | Green Fluo | Green Neutral | Red Fluo | Red Neutral | Red Dark Short Lived | Red Dark Long-Lived | Red Bleached |
|---|---|---|---|---|---|---|---|
| **Green Fluo** | 0 | 0 | 5e-7 | 0 | 0 | 0 | 0 |
| **Green Neutral** | 0 | 0 | 0 | 0.0001 | 0 | 0 | 0 |
| **Red Fluo** | 0 | 0 | 0 | 0 | 5e-6 | 1.5e-5 | 1e-5 |
| **Red Neutral** | 0 | 0 | 0 | 0 | 0 | 0 | 1e-5 |
| **Red Dark Short-Lived** | 0 | 0 | 0.0001<br>20* | 0 | 0 | 0 | 1e-5 |
| **Red Dark Long-Lived** | 0 | 0 | 0.005<br>0.001* | 0 | 0 | 0 | 1e-5 |

* Thermal exchange rates

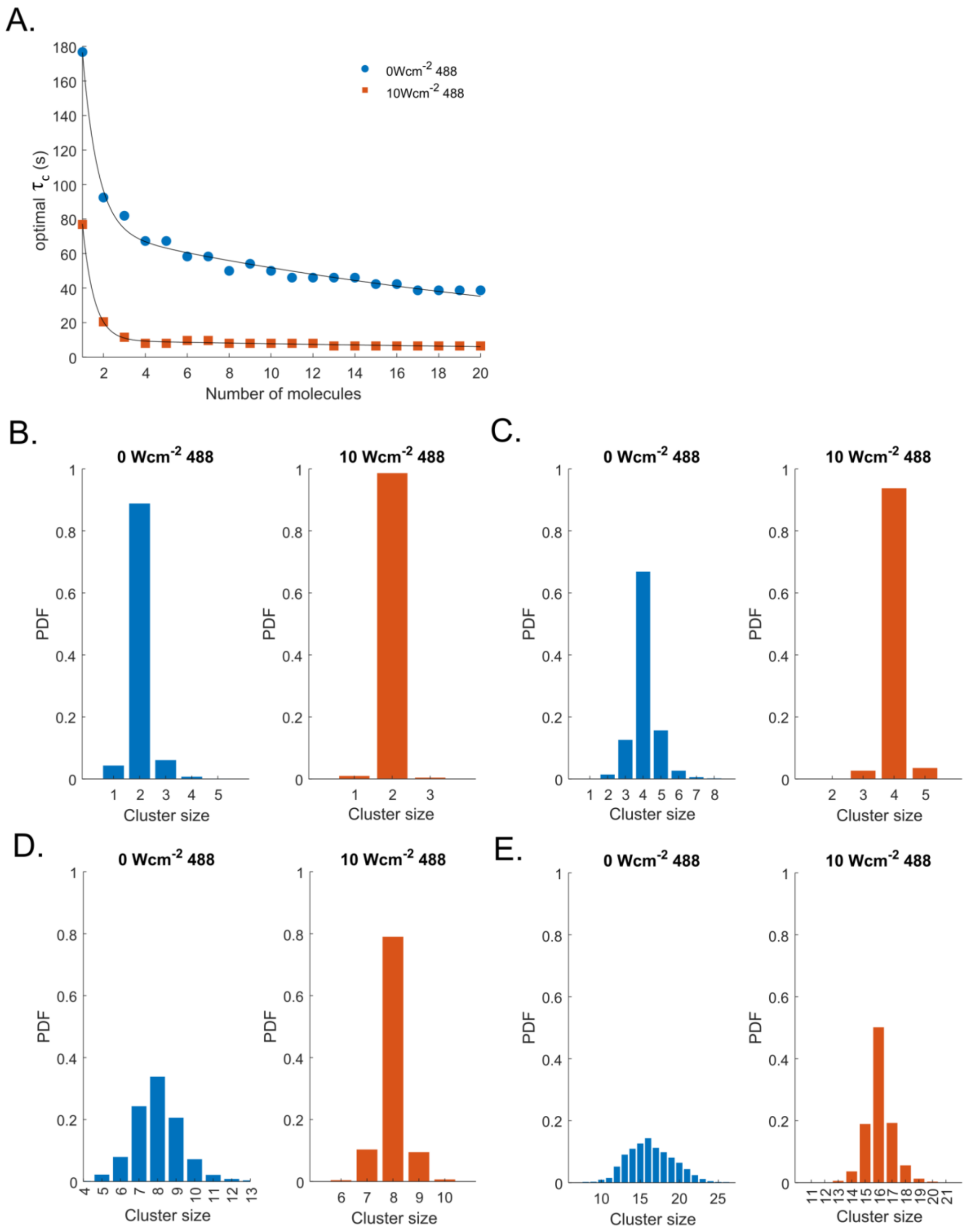


**Supplementary Figure S3. Counting of simulated data by off-time thresholding.** A) Optimal $\tau_c$ curves were calculated as described by Lee et al[5] from simulated monomers of mEos4b under illumination with 500W/cm$^2$ 561-nm, 1W/cm$^2$ 405-nm and 0 (blue circles) or 10W/cm$^2$ (orange squares) 488-nm light**.** Dimers (B), tetramers (C), octamers (D) and 16-mers (E) were reconstituted from simulated monomers with a labeling efficiency of 1 and counted by off-time thresholding using optimal $\tau_c$ values.[5]

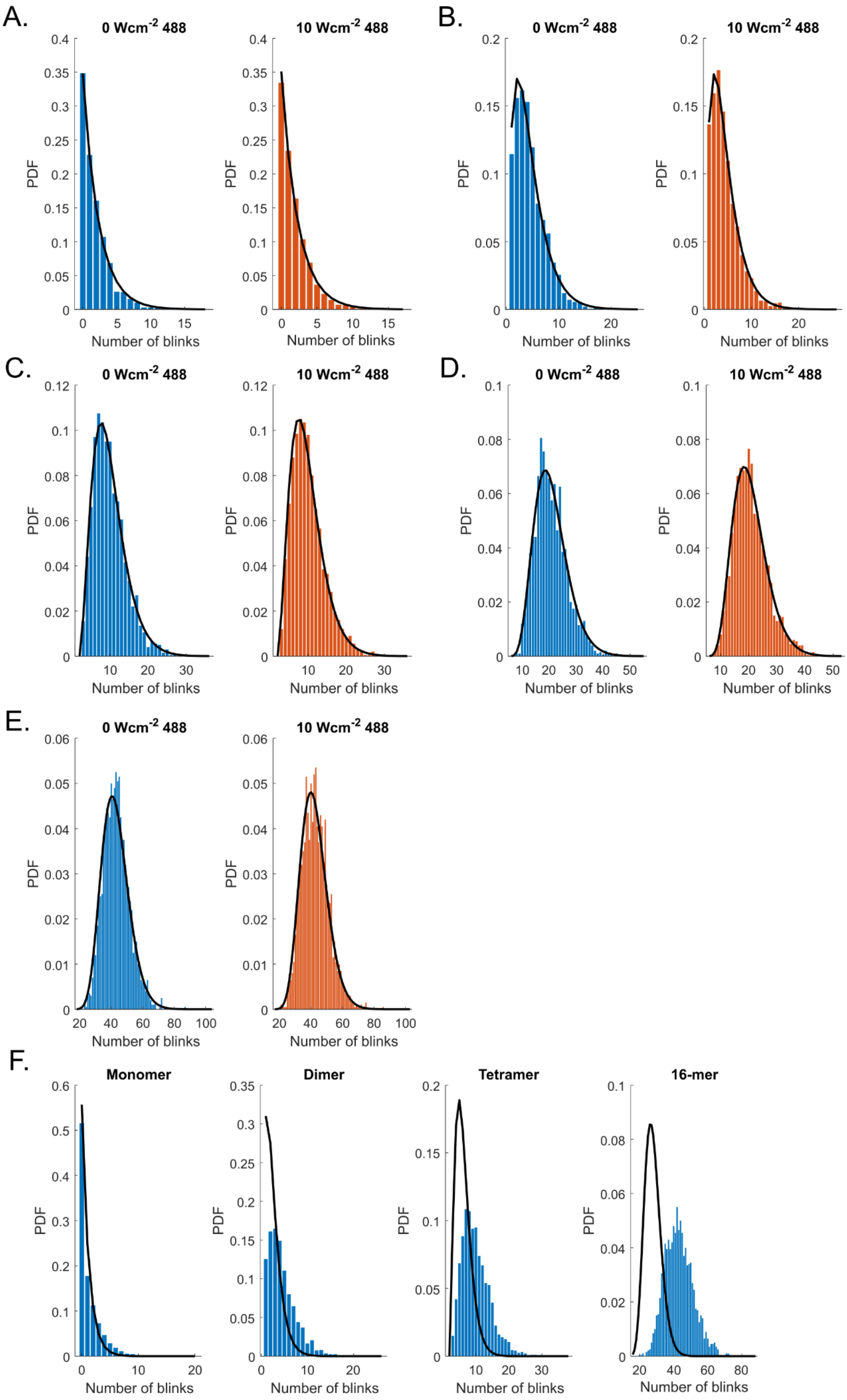
A.
0 Wcm-2 488
10 Wcm-2 488
B.
0 Wcm-2 488
10 Wcm-2 488
C.
0 Wcm-2 488
10 Wcm-2 488
D.
0 Wcm-2 488
10 Wcm-2 488
E.
0 Wcm-2 488
10 Wcm-2 488
F.
Monomer
Dimer
Tetramer
16-mer
PDF
Number of blinks

**Supplementary Figure S4. Counting of simulated data by blinking statistics** Monomers (A), dimers (B), tetramers (C), octamers (D) and 16-mers (E) were reconstituted with a labeling efficiency of 1 and histograms of the number of blinks (nBlink) were plotted. Solid lines indicate fits of the p-value (A) or fits of negative binomial distributions (B-E) according to ref [8], assuming a labeling efficiency of 1. F) A 20% error was added to the p-value, resulting in large deviations between the expected nBlink distribution (solid lines) and measured data (blue bars) at high stoichiometries.

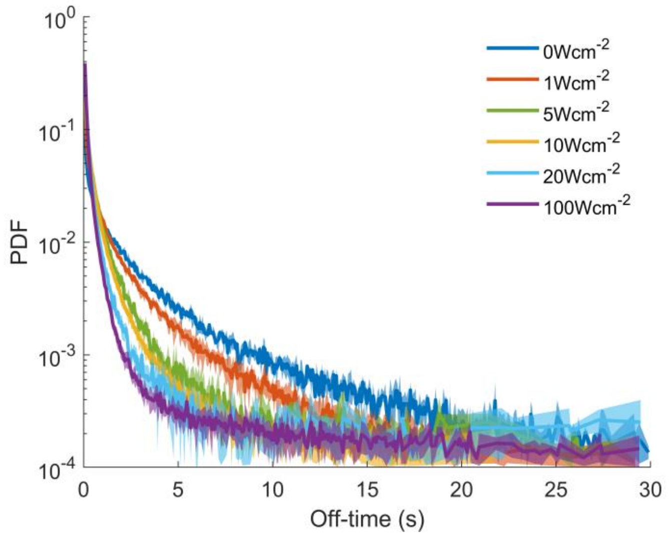


**Supplementary Figure S5. 405 illumination reduces the off-time duration of mEos4b.** Off-time histograms of mEos4b molecules embedded in PAA under alternating illumination with 500W/cm$^2$ 561-nm light (70 ms) and 1-100 W/cm$^2$ 405-nm (8.2 ms). Data represent mean ± s.d. of ≥ 3 measurements.

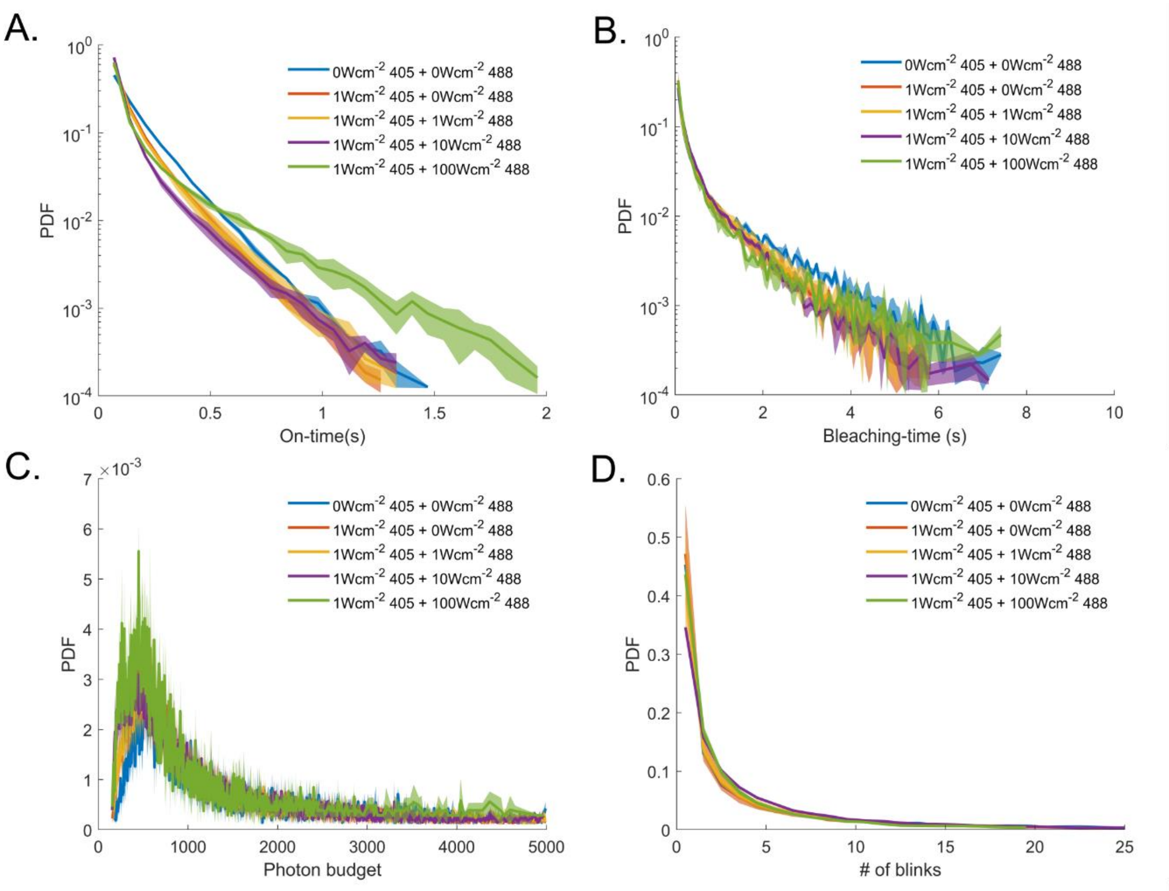


**Supplementary Figure S6. Effects of 488-nm light illumination on the red-state of mEos4b.** On-time histograms (A), bleaching-time histograms (B), photon-budget histograms (C), and nBlink histograms (D) of mEos4b molecules embedded in PAA under alternating illumination with 500W/cm$^2$ 561-nm light (70 ms) and 1W/cm$^2$ 405-nm plus 0-100 W/cm$^2$ 488-nm light (8.2 ms). Data represent mean ± s.d. of ≥ 3 measurements.

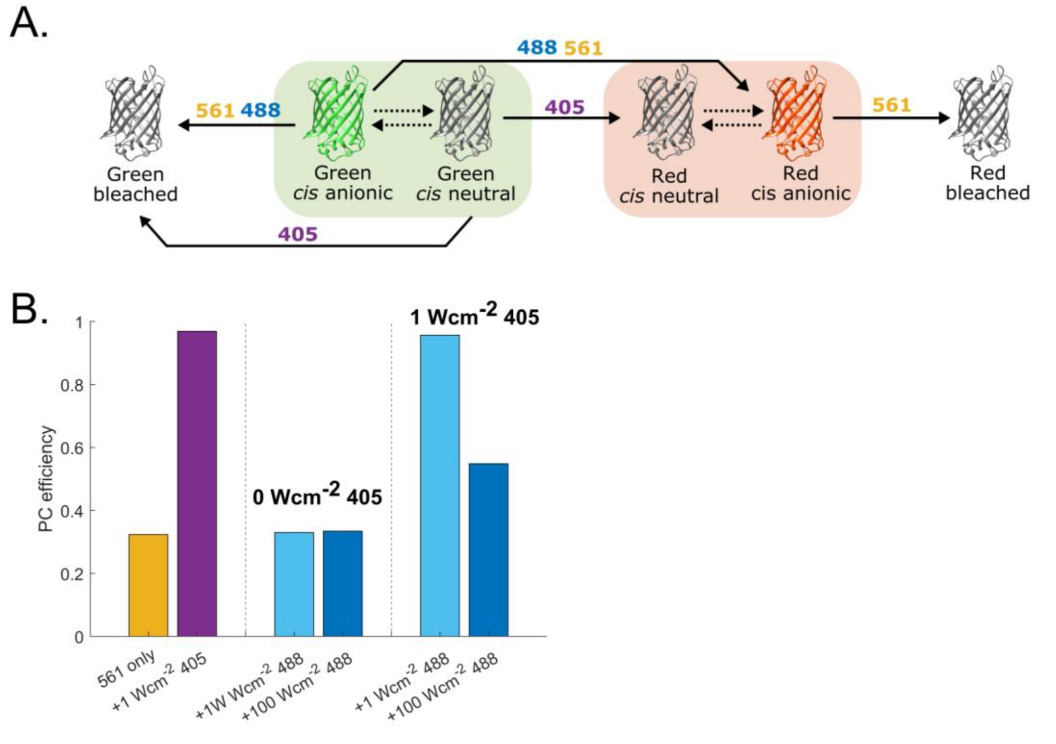


**Supplementary Figure S7. Simulations of the effects of 488-nm and 405-nm light on the photoconversion efficiency of mEos4b**. A) Simplified photophysical model of mEos4b to illustrate the process of photoconversion and bleaching under 405, 488 and 561-nm light illumination. Note that the simulations have been performed with the complete photophysical model of mEos4b (Supplementary Figure 1, Supplementary Note 2). B) PCE of simulated mEos4b molecules under illumination with 500W/cm$^2$ 561-nm light, 0-1 W/cm$^2$ 405-nm light and 0-10W/cm$^2$ 488-nm light.

**Supplementary Table S2. Phototransformation quantum yields and thermal exchange rates used for the ensemble simulations of mEos4b.**

| | **Green Fluo** | **Green Neutral** | **Green dark Short-Lived** | **Green dark Long-Lived** | **Green Bleached** | **Red Fluo** | **Red Neutral** | **Red Dark Short Lived** | **Red Dark Long-Lived** | **Red Bleached** |
|---|---|---|---|---|---|---|---|---|---|---|
| **Green Fluo** | 0 | 0 | 3.5e-5 | 5e-5 | 2.5e-6 | 5e-7 | 0 | 0 | 0 | 0 |
| **Green Neutral** | 0 | 0 | 0 | 0 | 2.5e-6 | 0 | 0.0001 | 0 | 0 | 0 |
| **Green dark Short-Lived** | 0.001<br>0.1* | 0 | 0 | 0 | 2.5e-6 | 0 | 0 | 0 | 0 | 0 |
| **Green dark Long-Lived** | 0.01<br>0.001* | 0 | 0 | 0 | 2.5e-6 | 0 | 0 | 0 | 0 | 0 |
| **Red Fluo** | 0 | 0 | 0 | 0 | 0 | 0 | 0 | 5e-6 | 1.5e-5 | 1e-5 |
| **Red Neutral** | 0 | 0 | 0 | 0 | 0 | 0 | 0 | 0 | 0 | 1e-5 |
| **Red Dark Short-Lived** | 0 | 0 | 0 | 0 | 0 | 0.0001<br>20* | 0 | 0 | 0 | 1e-5 |
| **Red Dark Long-Lived** | 0 | 0 | 0 | 0 | 0 | 0.005<br>0.001* | 0 | 0 | 0 | 1e-5 |

* Thermal exchange rates

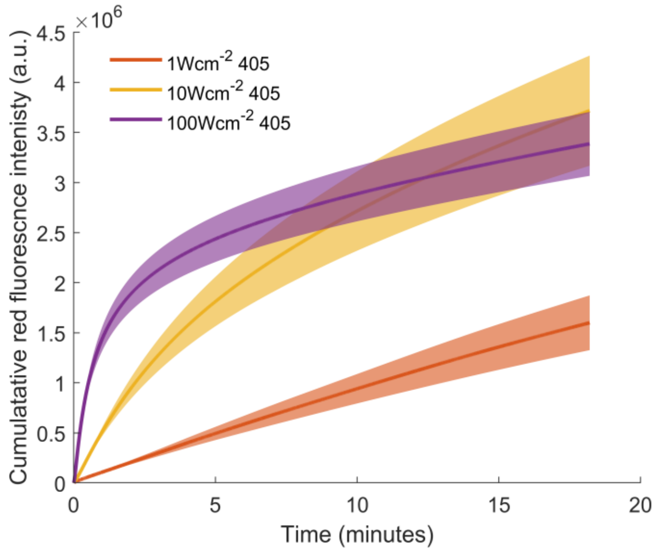


**Supplementary Figure S8. Ensemble level measurements of mEos4b PCE.** The cumulative red intensity of mEos4b molecules embedded in PAA under alternating illumination with 500W/cm$^2$ 561-nm light (70 ms) and 1-100W/cm$^2$ 405-nm light (8.2 ms). The cumulative red intensity can be used as a measure for the PCE because the red-state photon budget is not dependent on the 405-nm light intensity under PALM conditions (Supplementary Figure 9). Data represent mean ± s.d. of 2 measurements.

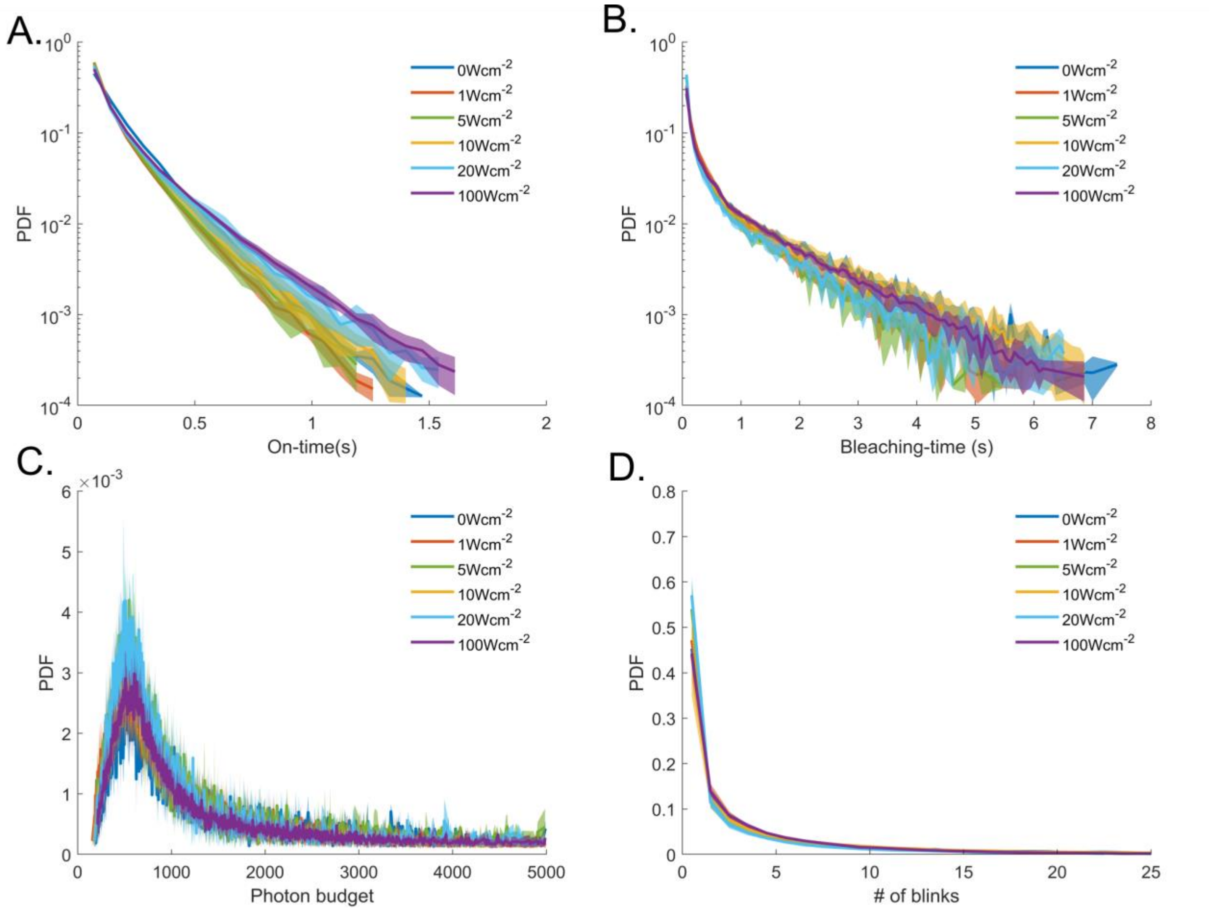


**Supplementary Figure S9. Effects of 405 illumination on the red-state of mEos4b.** On-time histograms (A), bleaching-time histograms (B), photon budget histograms (C), and nBlink histograms (D) of mEos4b molecules embedded in PAA under alternating illumination with 500W/cm$^2$ 561-nm light (70 ms) and 1-100W/cm$^2$ 405-nm light (8.2 ms). Data represent mean ± s.d. of ≥ 3 measurements.

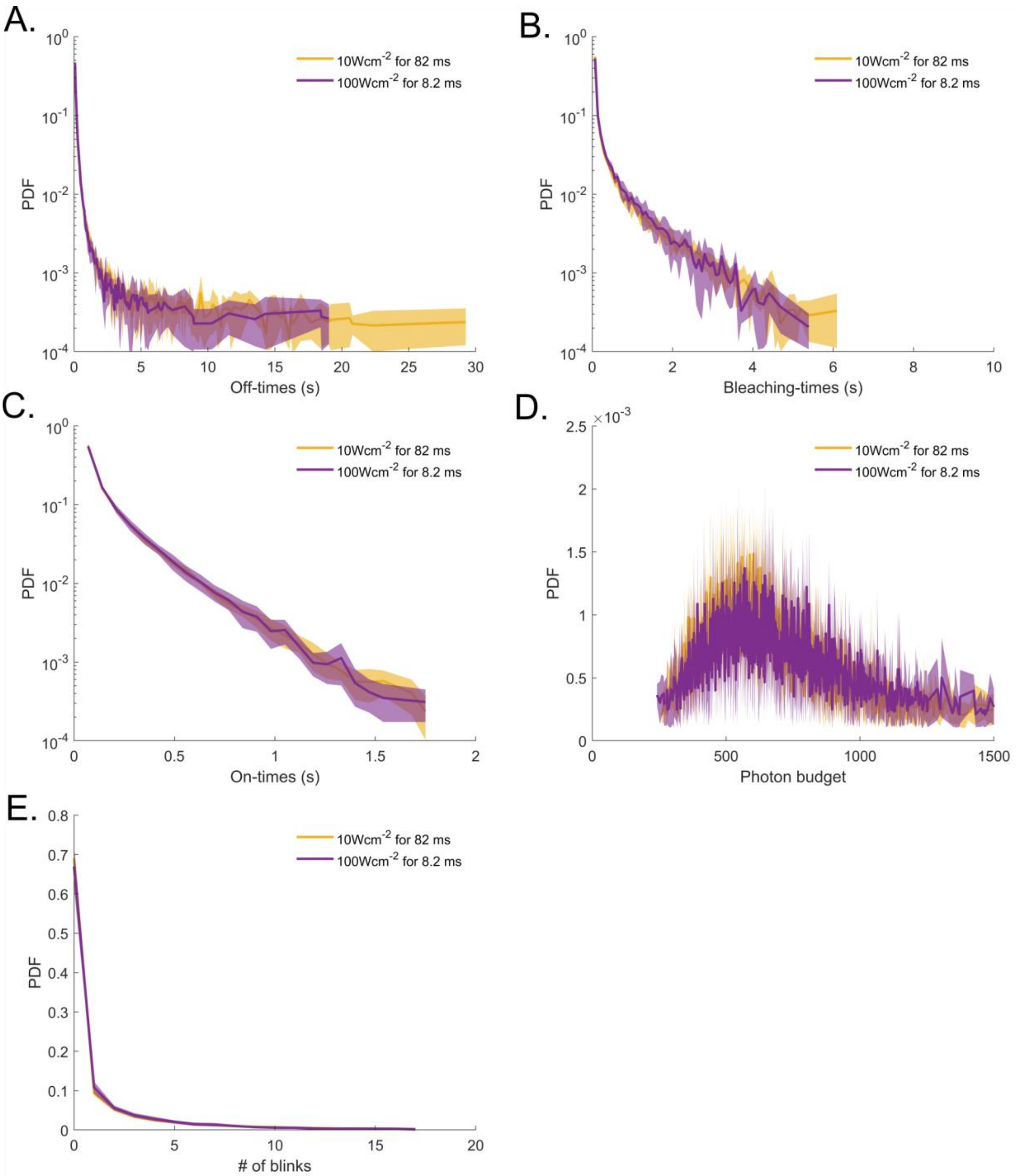


**Supplementary Figure S10. Effects of 405-nm light intensity on the red-state of mEos4b.** Off-time histograms (A), bleaching-time histograms (B), on-time histograms (C), photon-budget histograms (D), and nBlink histograms (E) of mEos4b molecules embedded in PAA under alternating illumination with 500W/cm$^2$ 561-nm light (70 ms) and 10 or 100W/cm$^2$ 405-nm light for 82 and 8.2 ms respectively. Data represent mean ± s.d. of ≥ 3 measurements.

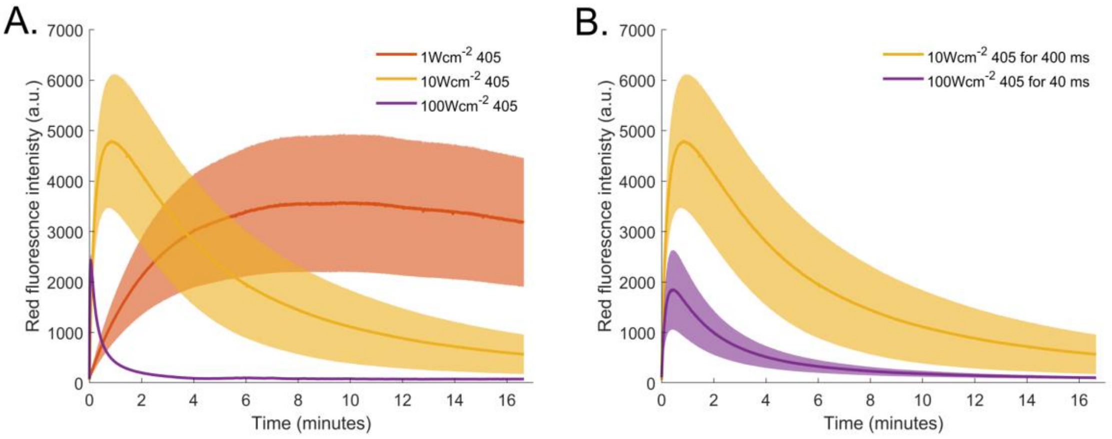


**Supplementary Figure S11. Effect of 405-nm illumination intensity on the PCE of mEos3.2.** Ensemble fluorescent time traces of mEos3.2 molecules embedded in PAA. The red fluorescent intensity was measured every 500 ms by illumination with 8 W/cm$^2$ 561-nm light for 20 ms. A) Samples were exposed to 400 ms pulses of 405-nm light (1-100W/cm$^2$). B) Samples were exposed to 40 and 400 ms pulses of 100 and 10W/cm$^2$ 405-nm respectively. Data represent mean ± s.d. of 3 measuments.

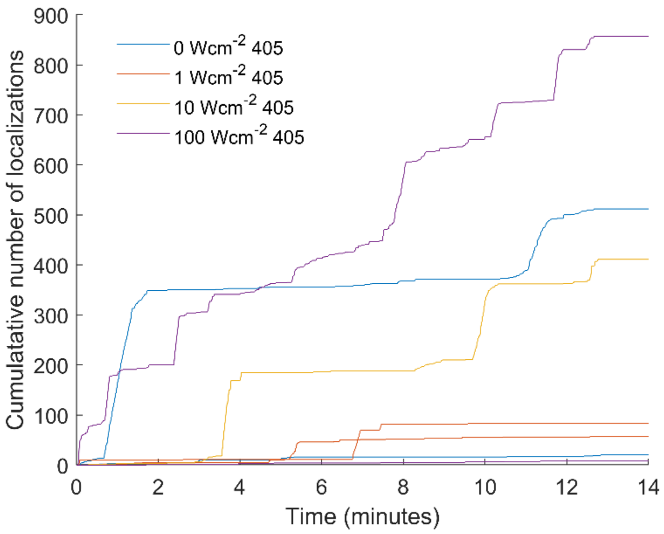


**Supplementary Figure S12. Red fluorescent impurities in the PAA samples.** The cumulative number of localizations detected on PAA samples without mEos4b molecules under alternating illumination with 500W/cm$^2$ 561-nm light (70 ms) and 0-100W/cm$^2$ 405-nm light (8.2 ms). Compared to the typical samples containing mEos4b (> 20000 clusters of localizations), the number of detected impurities is negligible. Of note, this plot shows the raw number of detections, while in the process of clustering, suspicious detections are removed so the number of spurious localizations contributing to the final data is even lower. Importantly, although there is variation in the number of detected impurities between measurements, there is no correlation to the 405-nm light intensity.

### Supplementary Notes

#### Supplementary Note 1: evaluation of counting approaches from simulated data

To count the simulated data by off-time thresholding, we calculated the optimal off-time threshold ($\tau_c$), which is dependent on the time between the photoconversion of different molecules, the off-time durations and the stoichiometry of the complex of interest, as described by Lee et al (2012).[5] Interestingly, the optimal $\tau_c$ value (Supplementary Figure 3A) was lower for the data simulated under illumination with 488-nm light. Since the PC kinetics (i.e. the average time between the photoconversion of different molecules) and the red-state bleaching-times (Supplementary Figure 2) were not significantly changed by the addition of 488-nm light, it can be concluded that this decrease in optimal $\tau_c$ reflects the shortened off-time durations. Using these optimal $\tau_c$ values to count the reconstituted oligomers of different sizes, the simulated data under 488-nm illumination showed a much smaller distribution in retrieved stoichiometries, resulting from the reduced trace mixing consecutive to the shorter off-time durations (Supplementary Figures S2C and S3). It should be noted however that the mean retrieved stoichiometry was always correct independently from the width of the distributions (Supplementary Figure S3). Yet, under experimental conditions, including reduced PCE and other sources of noise, we expect that reduction of the red-state off-times will facilitate counting.

In contrast to counting by off-time thresholding, counting by assessing the number of blinks per cluster appeared insensitive to the addition of 488-nm light. This is not surprising, as the number of blinks was essentially not changed (Supplementary Figures S4 and S2) and this method, at least in theory, is not sensitive to the duration of the off-times. However, the simulations highlighted an important caveat: we noticed that errors in the fit of the bleaching probability (p-value)[6] based on the nBlink histogram of the monomer data (which is typically affected by e.g. impurities or residual localization errors), translate into substantial errors when aiming to model large oligomers (Supplementary Figure S4F). Thus we suspect that counting by nBlink statistics may be inappropriate to investigate stoichiometries of large complexes.

#### Supplementary Note 2: Effects of 405-nm and 488-nm light on the PCE of mEos4b according to current photophysical model of mEos4b as proposed by Thedie et al.[7]

The PCE is determined by the ratio between the bleaching and PC quantum yields, which are assumed to be wavelength independent. In the absence of 405-nm illumination, residual photoconversion (PC) is induced from the *cis*-anionic state of the chromophore, which absorbs 561-nm and 488-nm light. Thus, in the absence of 405-nm light, addition of 488-nm light does not change the PCE as it increases the PC and bleaching rate in an equal manner (Supplementary Figure S7B). On the other hand, 405-nm light is absorbed by the *cis*-neutral state of the chromophore, for which the ratio in quantum yield between bleaching and PC is much more in favor of PC compared to the *cis*-anionic state. As a result, addition of 405-nm light increases the PCE as there is less bleaching from the *cis*-anionic state by 561-nm light, according to the model presented in Supplementary Figure S7A (Supplementary Figure S7B). Finally, in the situation where there is both 405-nm and 488-nm illumination, increasing the 488-nm intensity decreases the PC efficiency because the overall bleaching rate will increase more than the overall PC rate, which is mainly determined by the 405-nm intensity (Supplementary Figure S7B).

**Supplementary Note 3: mEos4b red-state photobleaching by 405-nm light**

The ensemble level data (Figure 3) show that 405-nm light decreases the red-state fluorescence of mEos4b in a dose dependent manner, which seems in contradiction with the PALM data (Supplementary Figure S9) showing that increasing the 405-nm light intensity does not increase red-state bleaching under PALM conditions. This apparent contradiction may be explained by the massive bleaching of the red-state by the intense 561-nm light under PALM conditions, hiding the contribution of the 405-nm light.

**Supplementary Note 4: Use of in vitro samples and non-quantitative evaluation of the PCE**

The use of *in vitro* samples allowed us to avoid any bias due to potentially uncontrolled parameters associated to real oligomeric structures such as multiple oligomeric states of unknown fractional populations; polarization dependent effects due to preferred orientations of the fluorophores linked to a multimeric target, notably in fixed cells; and possible energy transfer between fluorophores decorating such a target. However, a shortcoming of our approach is that the complex photophysical behavior of mEos4b and other PCFPs may differ from that in the specific environments of biological samples.[3,9] Our studies could benefit from more advanced *in vitro* immobilization methods such as those recently introduced, which in principle allow to apply environmental changes to the sample.[10,11]

Importantly, we refrained from extracting quantitative values for PCEs for the following reason: we observed that the cumulative curves of photoconverted red molecules never reached a flat plateau, especially at low 405 nm intensity. We verified that our samples were essentially devoid of impurities that could cause the observed slow rise (Supplementary Figure 12). Thus, we suspect that this could originate from the formation of ultra-long-lived dark states in red mEos4b that would induce very slow reappearance of red fluorescence. Another reason could be the anisotropic behavior of some of the fluorophores, as free tumbling of mEos4b could be partially restricted in PAA. Under such conditions, a quantitative evaluation of PCE values should be based on fitting of the cumulative curves by a complete photophysical model. However, our attempts for such fitting gave unsatisfactory results, because such complete model is still not available and because other experimental factors such as heterogeneity of the used lasers through the field of view or residual sample drift cannot be easily modeled.

**Supplementary Note 5: Estimation of the lifetime of the intermediate involved in nonlinear photobleaching by 405-nm light.**

We estimate the lifetime of the intermediate state with the assumption that non-linear bleaching becomes highly significant at a 405-nm power density of 100 W/cm² (Figure 3), although the process becomes noticeable at lower densities. The lifetime of the intermediate state should thus be long enough to allow absorption of a sufficient number of 405-nm photons so that photobleaching occurs under 100W/cm² 405-nm light illumination. Assuming that the mEos4b molecules are rapidly tumbling, the excitation rate (s-1) is given by:[3]

$$k = \varepsilon P \lambda \frac{(10^{-6}) Ln(10)}{\mathrm{N}_A hc}$$

where $\varepsilon$ [$M^{-1}cm^{-1}$] is the extinction coefficient of the considered species at wavelength $\lambda$ [nm]; $P$ [$W/cm^2$] is the laser power density; $N_A$ is the Avogadro number; $h$ is the Plank constant and $c$ is the speed of light. Estimating an extinction coefficient of 10 000 $M^{-1}cm^{-1}$ at maximum (based on the apparently weak absorption at 405 nm that we noticed in ref [12]), the excitation rate of the intermediate under 100W/cm² 405-nm light illumination is 7.7 $10^4$ $s^{-1}$. Furthermore, assuming a strong photobleaching quantum yield of $10^{-3}$, it would take 1000 absorption events to effectively produce photobleaching. Thus we estimate the lifetime of the intermediate to be at minimum ~15 ms, a value at least one order of magnitude larger than the estimated triplet state lifetime.[13]

### Supplementary bibliography


(1) De Zitter, E.; Ridard, J.; Thédié, D.; Adam, V.; Lévy, B.; Byrdin, M.; Gotthard, G.; Van Meervelt, L.; Dedecker, P.; Demachy, I.; Bourgeois, D. Mechanistic Investigations of Green MEos4b Reveal a Dynamic Long-Lived Dark State. *J. Am. Chem. Soc.* **2020**, *142* (25), 10978–10988. https://doi.org/10.1021/jacs.0c01880.

(2) Ovesný, M.; Křížek, P.; Borkovec, J.; Švindrych, Z.; Hagen, G. M. ThunderSTORM: A Comprehensive ImageJ Plug-in for PALM and STORM Data Analysis and Super-Resolution Imaging. *Bioinformatics* **2014**, *30* (16), 2389–2390. https://doi.org/10.1093/bioinformatics/btu202.

(3) Avilov, S.; Berardozzi, R.; Gunewardene, M. S.; Adam, V.; Hess, S. T.; Bourgeois, D. In Cellulo Evaluation of Phototransformation Quantum Yields in Fluorescent Proteins Used As Markers for Single-Molecule Localization Microscopy. *PLoS ONE* **2014**, *9* (6), e98362. https://doi.org/10.1371/journal.pone.0098362.

(4) De Zitter, E.; Thédié, D.; Mönkemöller, V.; Hugelier, S.; Beaudouin, J.; Adam, V.; Byrdin, M.; Van Meervelt, L.; Dedecker, P.; Bourgeois, D. Mechanistic Investigation of MEos4b Reveals a Strategy to Reduce Track Interruptions in SptPALM. *Nat. Methods* **2019**, *16* (8), 707–710. https://doi.org/10.1038/s41592-019-0462-3.

(5) Lee, S.-H.; Shin, J. Y.; Lee, A.; Bustamante, C. Counting Single Photoactivatable Fluorescent Molecules by Photoactivated Localization Microscopy (PALM). *Proc. Natl. Acad. Sci.* **2012**, *109* (43), 17436–17441. https://doi.org/10.1073/pnas.1215175109.

(6) Fricke, F.; Beaudouin, J.; Eils, R.; Heilemann, M. One, Two or Three? Probing the Stoichiometry of Membrane Proteins by Single-Molecule Localization Microscopy. *Sci. Rep.* **2015**, *5* (1), 14072. https://doi.org/10.1038/srep14072.

(7) Thédié, D.; Berardozzi, R.; Adam, V.; Bourgeois, D. Photoswitching of Green MEos2 by Intense 561 Nm Light Perturbs Efficient Green-to-Red Photoconversion in Localization Microscopy. *J. Phys. Chem. Lett.* **2017**, *8* (18), 4424–4430. https://doi.org/10.1021/acs.jpclett.7b01701.

(8) Baldering, T. N.; Bullerjahn, J. T.; Hummer, G.; Heilemann, M.; Malkusch, S. Molecule Counts in Complex Oligomers with Single-Molecule Localization Microscopy. *J. Phys. Appl. Phys.* **2019**, *52* (47), 474002. https://doi.org/10.1088/1361-6463/ab3b65.

(9) Sun, M.; Hu, K.; Bewersdorf, J.; Pollard, T. D. Sample Preparation and Imaging Conditions Affect MEos3.2 Photophysics in Fission Yeast Cells. *Biophys. J.* **2021**, *120* (1), 21–34. https://doi.org/10.1016/j.bpj.2020.11.006.

(10) Platzer, R.; Rossboth, B. K.; Schneider, M. C.; Sevcsik, E.; Baumgart, F.; Stockinger, H.; Schütz, G. J.; Huppa, J. B.; Brameshuber, M. Unscrambling Fluorophore Blinking for Comprehensive Cluster Detection via Photoactivated Localization Microscopy. *Nat. Commun.* **2020**, *11* (1), 4993. https://doi.org/10.1038/s41467-020-18726-9.

(11) Jusuk, I.; Vietz, C.; Raab, M.; Dammeyer, T.; Tinnefeld, P. Super-Resolution Imaging Conditions for Enhanced Yellow Fluorescent Protein (EYFP) Demonstrated on DNA Origami Nanorulers. *Sci. Rep.* **2015**, *5* (1), 14075. https://doi.org/10.1038/srep14075.

(12) Adam, V.; Carpentier, P.; Violot, S.; Lelimousin, M.; Darnault, C.; Nienhaus, G. U.; Bourgeois, D. Structural Basis of X-Ray-Induced Transient Photobleaching in a Photoactivatable Green Fluorescent Protein. *J. Am. Chem. Soc.* **2009**, *131* (50), 18063–18065. https://doi.org/10.1021/ja907296v.
(13) Byrdin, M.; Duan, C.; Bourgeois, D.; Brettel, K. A Long-Lived Triplet State Is the Entrance Gateway to Oxidative Photochemistry in Green Fluorescent Proteins. *J. Am. Chem. Soc.* **2018**, *140* (8), 2897–2905. https://doi.org/10.1021/jacs.7b12755.
